# Deep learning-guided identification of bacteriophage receptor-binding protein candidates for foodborne pathogen detection

**DOI:** 10.64898/2026.08.03.742407

**Authors:** Danitza Xiomara Romero-Calle, Cristhian Carrasco, Bilal Javed, Elena-Alexandra Alexa

## Abstract

Foodborne pathogens including *Salmonella* spp., *Escherichia coli* and *Listeria monocytogenes* cause an estimated 600 million illnesses annually. Yet conventional detection methods remain slow, costly, or insufficiently specific for routine food safety surveillance. Phage receptor-binding proteins (RBPs) are attractive recognition elements for biosensors, but their extensive sequence diversity limits reliable computational identification.

Here, we present a systematic open-source computational pipeline for identifying and structurally characterising high-confidence RBP candidates from phage genomes targeting these three priority pathogens. The pipeline integrates four stages: deep learning-based RBP prediction, protein structure prediction, structural homology validation, and exploratory molecular docking.

Applied to a quality-controlled dataset of 247 complete phage genomes retrieved from the National Center for Biotechnology Information Nucleotide database (31,752 total protein sequences), PhageRBPdetect, built on the ESM-2 protein language model, identified 653 high-confidence RBP candidates. Foldseek structural homology validation against PDB100 confirmed 13 candidates with a structural match probability of 1.0 to known phage adsorption proteins, spanning four structural archetypes. ESMFold-predicted structures showed strong confidence, with a mean model confidence score of 0.89 and a 90.2% prediction success rate. Exploratory rigid-body docking identified YDV08491.1, an E. coli-targeting candidate, as having the most energetically favourable predicted interaction, with a predicted binding energy of −132.3 kcal/mol against OmpF, supporting experimental prioritisation.

These candidates’ structural diversity and predicted host specificity support their future development as phage-based biosensors and biocontrol tools, and this reproducible, accessible pipeline offers a transferable strategy for prioritising RBP candidates in downstream functional studies, pending experimental validation.

**Author Summary:** Foodborne bacterial infections cause an estimated 600 million illnesses every year worldwide, with *Salmonella*, *Escherichia coli*, and *Listeria monocytogenes* among the most significant contributors. Detecting these pathogens quickly and specifically in food supply chains remains a major challenge for global food safety. Bacteriophages viruses that infect bacteria use surface proteins called receptor-binding proteins (RBPs) to recognize their bacterial hosts with remarkable precision, making RBPs promising building blocks for pathogen-specific biosensors. However, the huge sequence diversity of RBPs across phage genomes has made it difficult to reliably identify good candidates computationally.

We built an open-source pipeline that combines deep learning, protein structure prediction, and molecular docking to screen phage genomes for high-confidence RBP candidates. Applied to 247 phage genomes targeting the three pathogens above, our pipeline flagged 653 candidate RBPs, of which 13 showed strong structural similarity to known phage adsorption proteins. One candidate, targeting *E. coli*, showed a particularly strong predicted binding interaction with a bacterial surface protein, making it a priority target for experimental testing.

This pipeline gives researchers a reproducible, publicly available strategy for prioritizing phage proteins toward the development of biosensors and biocontrol tools for food safety.

## Introduction

Foodborne diseases caused by bacterial pathogens represent a major and persistent global public health burden [1, 2]. Among the most clinically and economically significant agents, *Salmonella* spp., *Escherichia coli,* and *Listeria monocytogenes* collectively account for a substantial proportion of these cases across all income settings [2]. Bacterial contamination of food causes approximately 600 million cases of foodborne illness and 420,000 annual deaths worldwide [1] This burden is disproportionately high in low- and middle-income countries, where surveillance infrastructure and rapid diagnostic capacity remain limited. The European Food Safety Authority and the European Centre for Disease Prevention and Control report that *Salmonella* and *Listeria monocytogenes* remain among the most significant zoonotic pathogens, with thousands of confirmed human cases annually [2].

Bacteriophages have shown extraordinary host specificity, determined primarily by the interaction between their receptor-binding proteins (RBPs) and bacterial surface receptors, emerging as compelling alternatives to conventional antibiotics with demonstrated efficacy across food safety, clinical, and environmental contexts [3, 4]. Central to this specificity are receptor-binding proteins (RBPs): structural components located at the phage tail tip that mediate the initial, irreversible recognition of host surface receptors. RBPs which include tail fiber proteins, tail spike proteins, and baseplate hub components determine phage host range with molecular precision, making them primary determinants of phage–host interaction specificity [5, 6]. Beyond their foundational role in phage biology, RBPs are increasingly being explored as components of biosensing and targeted antimicrobial strategies due to their highly specific host-recognition properties, particularly in complex food matrices where detection specificity is critical [7].

Despite their importance, the systematic identification and characterisation of RBPs from phage genomes remains technically challenging, primarily due to their extreme sequence diversity, low sequence conservation across phage families, and the absence of universal domain signatures that would allow reliable annotation using conventional homology-based tools. RBPs exhibit extreme sequence diversity even among phages targeting the same bacterial host, and many are annotated as hypothetical proteins in public databases, rendering sequence homology-based approaches insufficient for comprehensive discovery of functional RBP candidates from phage genomic data [5, 8] This annotation gap is particularly pronounced for phages targeting Gram-positive pathogens such as *L. monocytogenes,* where the structural diversity of tail architectures further limits the effectiveness of conventional bioinformatic approaches. The recent development of deep learning-based protein structure prediction tools particularly ESMFold [9] and the ESM-2 protein language model has created new opportunities for structure-guided RBP identification that transcend sequence-level limitations.

To accelerate discovery, several computational tools have been developed specifically for RBP prediction. Notably, PhageRBPdetect leverages a fine-tuned ESM-2 protein language model to identify RBPs directly from genomic sequences [10]. This tool enables binary classification of phage proteins as RBP or non-RBP with high sensitivity, even for divergent sequences lacking recognisable domain architecture. Complementary structural search tools such as Foldseek [11, 12] enable rapid comparison of predicted structures against experimentally resolved protein databases, providing independent structural validation through comparison with crystallographically confirmed phage adsorption components, a critical step for distinguishing genuine RBP folds from prediction artifacts.

Here, we present a systematic, open-source computational pipeline for the identification and structural characterization of high-confidence RBP candidates from phage genomes targeting *Salmonella* spp., *Escherichia coli*, and *Listeria monocytogenes*. Starting from 247 complete phage genomes retrieved from NCBI, the extraction of 31,752 protein sequences, applied PhageRBPdetect to identify 653 high-confidence candidates, predicted and validated structures via ESMFold and Foldseek, and performed protein–protein docking against known host surface receptors. This pipeline was designed for reproducibility and accessibility, with all code and datasets made publicly available to support adoption by research groups operating under computational or infrastructural constraints.

## Results

### High-Confidence RBP Identification Across Foodborne Pathogens

Of the 260 initial genomes, 247 passed quality filtering, representing a 95.0% retention rate.

The deep learning pipeline applied to 31,752 phage proteins identified 699 RBP candidates across the three pathogen groups. Of these, 653 met the high-confidence threshold of >0.9, representing 93.4% of initial candidates and forming the core dataset for structural characterization. Detection rates were consistent between *Salmonella* and *Escherichia* phages at 2.6% each, suggesting comparable RBP representation in their respective phage datasets. In contrast, *Listeria* phages yielded a markedly lower detection rate of 0.5%, a finding that likely reflects the comparatively smaller and less annotated Listeria phage genomic landscape currently available in public repositories. This disparity highlights a broader gap in phage genomics data- bases for Gram-positive foodborne pathogens, a limitation with direct implications for biosensor development targeting *Listeria monocytogenes* in food safety contexts (Table 2).

**Table 1.**
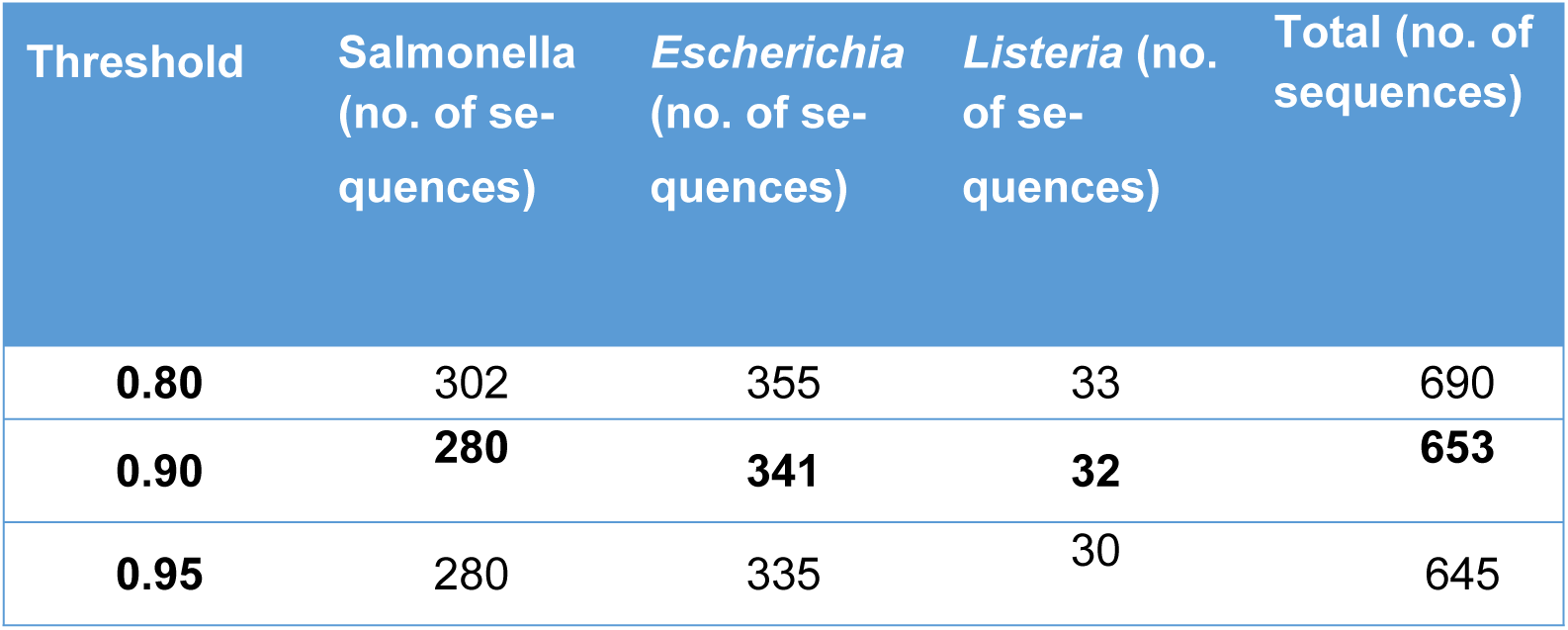
Sensitivity table in methods.

**Table 2.** Summary of phage RBP predictions across the three foodborne pathogen datasets.

| Pathogen | Total proteins | Predicted RBPs | RBP rate (%) | High-conf. RBPs (0.9 < score) | High-conf. rate (%) |
| --- | --- | --- | --- | --- | --- |
| ***Salmonella* phages** | 11,710 | 306 | %2.6 | 280 | %91.5 |
| ***Escherichia* phages** | 13,760 | 359 | %2.6 | 341 | %95 |
| ***Listeria* phages** | 6,282 | 34 | %0.5 | 32 | %94.1 |
| **Total** | 31,752 | 699 | %2.2 | 653 | %93.4 |

High-confidence RBPs: prediction score >0.9.

### Structural Quality Assessment of ESMFold Predictions

Of the 306 high-confidence RBP candidates submitted to ESMFold, 276 structures were successfully predicted, corresponding to a 90.2% success rate. Structural quality was evaluated using the predicted Local Distance Difference Test (pLDDT) score, where values above 0.85 were utilised as the threshold for reliable atomic-level predictions. The dataset showed strong overall confidence, with a mean pLDDT of 0.892 and a median of 0.904, suggesting that most predicted structures are suitable for downstream structural analyses. Consistent with this, 213 of 276 structures (77.2%) surpassed the pLDDT ≥0.85 threshold applied for candidate selection (Table 3). The 30 sequences for which prediction failed were predominantly identified as elongated tail fiber and spike proteins, based on the functional annotations present in the input FASTA headers. These failures occurred due to API time-out errors and request rejections, consistently correlated with sequences belonging to these long, structurally complex classes. This limitation is inherent to the remote execution of current AI-based protein folding tools on large or repetitive proteins, rather than specific to the dataset used here reinforcing the need for complementary validation of structurally complex RBP candidates in future work.

**Table 3.**
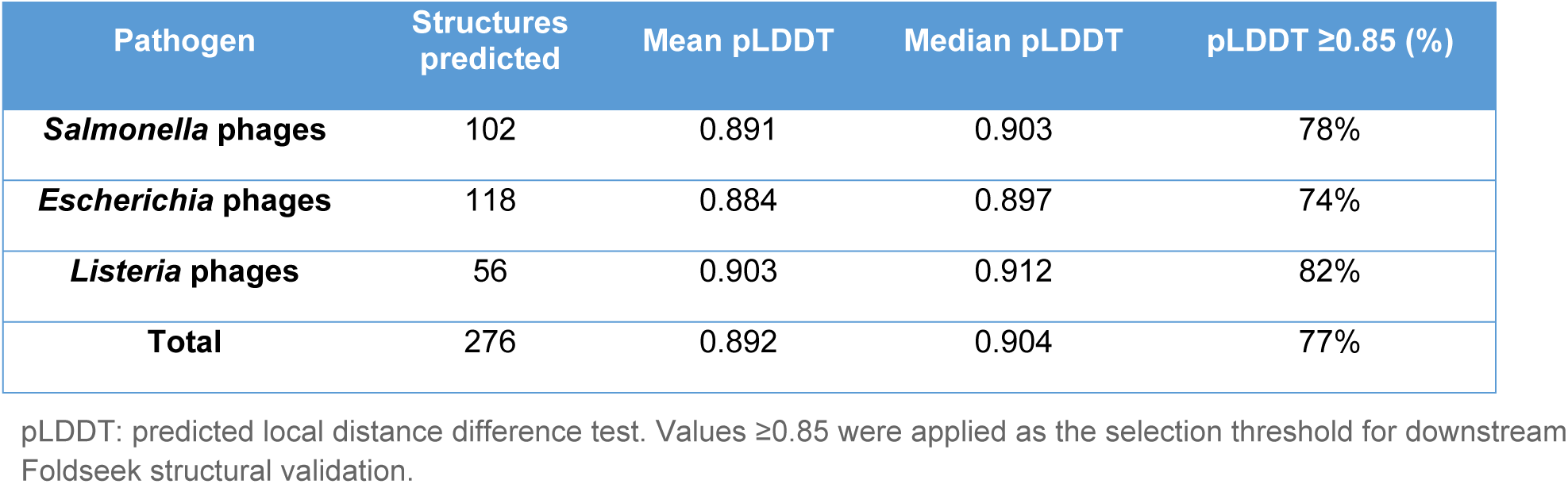
ESMFold structural prediction quality metrics by pathogen dataset.

### Structural Validation via Foldseek Against the PDB

To contextualize the predicted structures within the known phage proteome, the 15 top-ranked RBP candidates comprising the top 5 highest-scoring sequences based on prediction confidence scores from each of the three target pathogen groups (*Salmonella, Escherichia*, and *Listeria*) were submitted to structural homology search against PDB100. Thirteen of the 15 candidates (86.7%) returned statistically significant hits, defined by a probability ≥ 0.9 and E-value < 0.01, confirming structural correspondence with experimentally resolved phage proteins (Table 4).

**Table 4.**
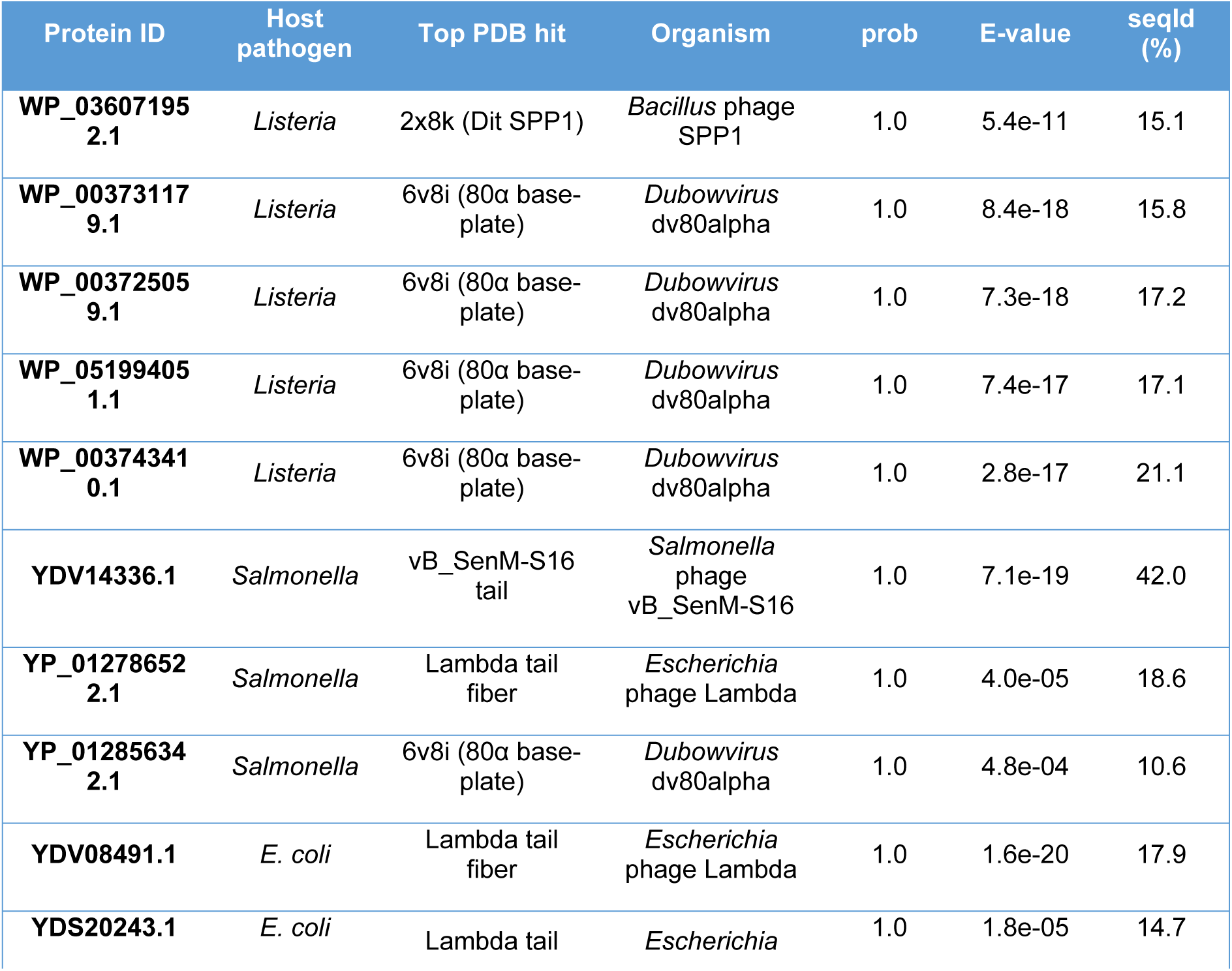
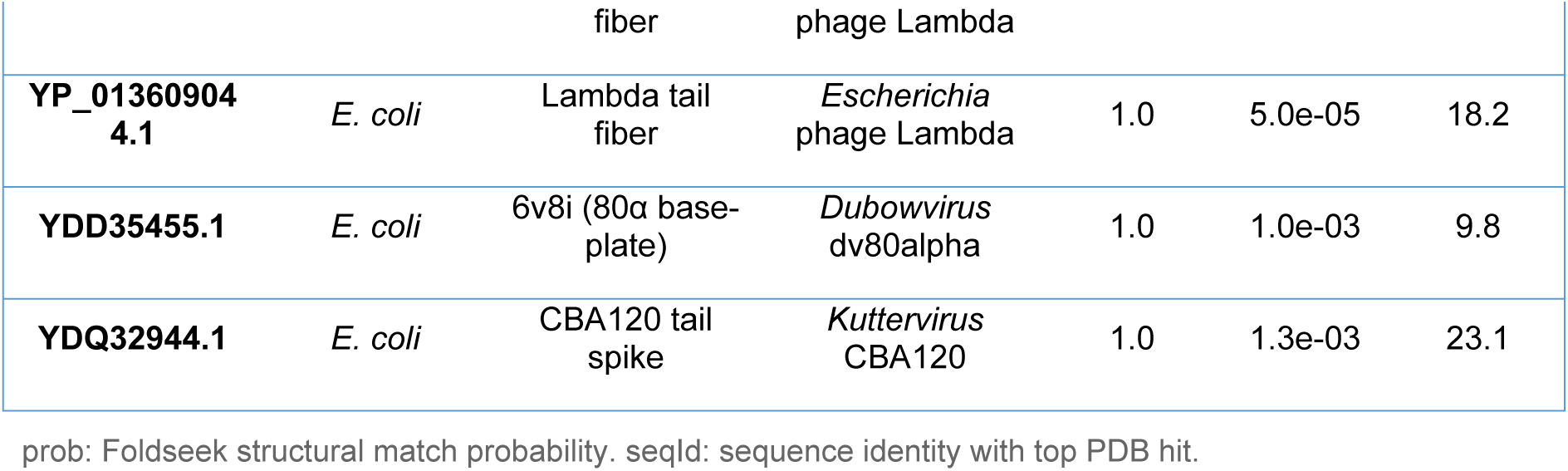
Foldseek structural validation of the 13 final RBP candidates against PDB100.

Four structural archetypes emerged from this analysis. The SPP1 Dit/80α baseplate hub was the most represented, identified in seven candidates spanning all three pathogen groups (five *Listeria-*, one *Salmonella-*, and one *Escherichia*-targeting), suggesting broad structural conservation of baseplate-mediated adsorption mechanisms across genera. The Lambda tail fiber archetype was identified in four candidates three *Escherichia coli*-targeting and one Salmonella-targeting reinforcing this cross-host pattern and providing a strong experimental foundation for downstream biosensor engineering, given that Lambda tail fiber is among the most structurally and functionally characterised RBP scaffolds in phage biology. In contrast, two archetypes showed strict host specificity: a single *Salmonella*-targeting candidate matched the vB_SenM-S16 tail protein at 42.0% sequence identity the highest identity observed in this analysis while a single *E. coli*-targeting candidate (YDQ32944.1) matched the CBA120 tail spike at 23.1% identity, the second-highest match overall. Both patterns are consistent with structural conservation despite moderate sequence divergence, increasingly documented in phage tail proteins [16].

### Protein–Protein Docking of Top RBP Candidates Against Host Surface Receptors

Protein–protein docking was performed to evaluate the binding potential of the top-ranked RBP candidate per pathogen group against their respective host surface receptors. Results revealed a spectrum of interaction energetics that reflect both the diversity of RBP binding mechanisms and the inherent limitations of rigid-body docking models for structurally complex interfaces (Table 5).

**Table 5.**
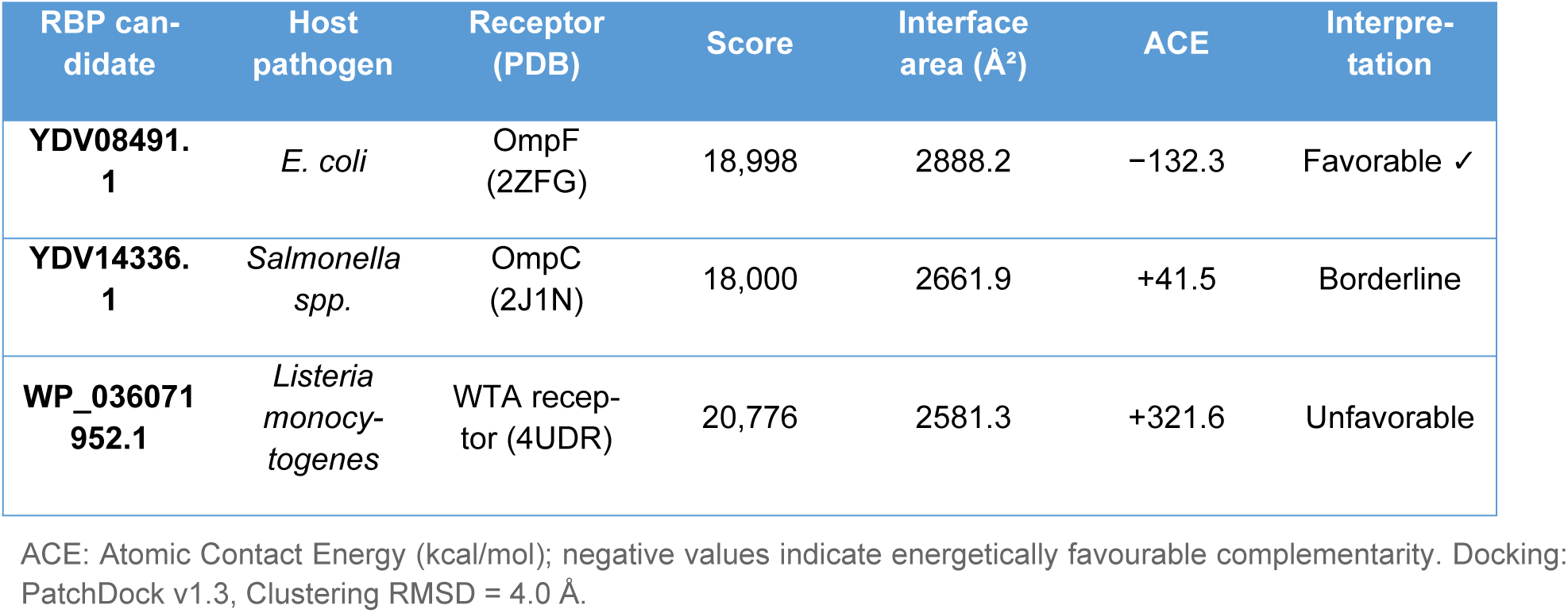
Protein–protein docking results for top RBP candidates against host surface receptors.

The most energetically favourable interaction was predicted for YDV08491.1 against OmpF from *Escherichia coli* (ACE = −132.3 kcal/mol; interface area = 2,888.2 Å²), suggesting a predicted binding interface with favourable geometric and energetic complementarity. While these results are exploratory and require experimental validation, YDV08491.1 emerges as the high-est-priority candidate for experimental binding assays among the three pathogen groups evaluated.

In contrast, YDV14336.1 against OmpC from *Salmonella* spp. yielded a borderline atomic contact energy (ACE = +41.5 kcal/mol) alongside a high geometric score of 18,000, suggesting favourable geometric complementarity despite suboptimal energetic parameters. This discrepancy may reflect conformational flexibility in the OmpC binding interface not captured by the rigid-body docking algorithm, a limitation that molecular dynamics simulations could address in future work [17].

Consequently, the least favourable result was obtained for WP_036071952.1 against the wall teichoic acid receptor of *Listeria monocytogenes* (ACE = +321.6 kcal/mol). Rather than indicating weak binding, this outcome most likely reflects a fundamental incompatibility between the protein-only docking model and the carbohydrate-dependent nature of WTA-mediated phage adsorption, a mechanism that requires glycan moieties not represented in standard PDB structures. This finding underscores the need for carbohydrate-aware docking approaches when modelling Listeria phage–host interactions, a gap that remains largely unaddressed in current computational pipelines.

## Discussion

This study presents a systematic, multi-stage computational pipeline for the identification and structural characterisation of high-confidence RBP candidates from phage genomes targeting *Salmonella* spp., *Escherichia coli*, and *Listeria monocytogenes*. By integrating deep learning-based RBP prediction, ESMFold structure prediction, Foldseek structural validation, and exploratory protein–protein docking within a fully open-source, reproducible framework, we identified 13 structurally validated RBP candidates, providing a computationally grounded, resource-accessible foundation for subsequent experimental characterisation and downstream biosensor probe engineering.

A central challenge in RBP identification is their high sequence diversity, which limits the reliability of conventional annotation approaches [5]. Using PhageRBPdetect on the ESM-2 backbone, 653 high-confidence candidates were recovered from 31,752 proteins at a threshold of 0.90, with detection rates of 2.6% for both *Salmonella* and *Escherichia* datasets. *Listeria* phages exhibited a markedly lower detection rate (0.5%), likely reflecting their underrepresentation in public databases and the structural divergence of their tail architectures [10]. To assess whether the 0.90 threshold was driving this outcome, a comparison was performed across thresholds of 0.80 and 0.95: total candidates shifted by only 6.5% (690 to 645), and *Listeria* candidates remained almost unchanged (33 to 30), supporting the robustness of the selected cutoff. The minimal fluctuation in *Listeria* candidate counts across these threshold variations supports the interpretation that the lower detection rate reflects an intrinsic biological and representational feature of the dataset, rather than an artefact of stringency choices.

Independent structural validation using Foldseek supported the RBP annotation of 13 of the 15 highest-ranking candidates (86.7%), all of which showed their highest structural similarity to experimentally characterised phage adsorption proteins. Four structural archetypes emerged from the validated dataset. The SPP1 Dit/80α baseplate hub was the most frequently recovered, identified in seven candidates spanning all three pathogen groups (five *Listeria*-, one *Salmonella*-, and one *Escherichia*-targeting). This widespread distribution suggests that key structural features involved in baseplate-mediated adsorption have been conserved across evolution despite diversification of host range. The recurrence of these archetypes across multiple pathogen groups is consistent with the deep evolutionary conservation of phage tail architectures [8] and underscores the utility of structure-guided approaches for RBP annotation in datasets where sequence-based methods fail.

Although AlphaFold-based modelling yielded highly confident structural predictions for the identified receptor-binding protein (RBP) candidates, a nuanced interpretation of these models is required. The per-residue confidence metric, pLDDT, is strictly a local indicator of structural accuracy and does not necessarily correlate with thermodynamic stability or reflect global do-main–domain orientations [11]. Because phage RBPs often exhibit modular arrangements with flexible hinge regions or extended coiled-coil architectures, a high pLDDT score within isolated functional domains does not guarantee the precision of their relative spatial positioning. To mitigate the inherent limitations of single-sequence folding algorithms when processing modular phage RBPs with flexible hinge regions or extended architectures, our structure-guided screening was systematically complemented by independent structural homology validation via Foldseek.

Preliminary docking analysis supported the predicted host-recognition function of the *E. coli*-targeting candidate YDV08491.1, which displayed favourable binding energetics against OmpF (ACE = −132.3 kcal/mol). OmpF is a well-characterised phage receptor exploited by numerous siphoviruses, including phage Lambda the closest structural homolog of YDV08491.1 identified via Foldseek. For the *Salmonella*- and *Listeria*-targeting candidates, borderline or unfavourable docking outcomes likely reflect the limitations of rigid-body docking against static receptor structures rather than an absence of binding potential, consistent with previous observations on the inadequacy of protein-only models for carbohydrate-mediated adsorption interfaces [18].

These 13 Foldseek-validated candidates represent structurally characterised, computationally prioritised foundations for immediate experimental validation. The identification of YDV14336.1, with direct structural and sequence homology (42% identity) to a crystallographically resolved *Salmonella*-specific adsorption protein, provides a rational basis for recombinant expression and functional binding assays without requiring *de novo* structural characterisation. For biosensor development, these candidates require experimental verification of their host-binding kinetics prior to integration into diagnostic detection platforms.

Four primary limitations of this framework warrant explicit acknowledgment. First, all analyses are computational, and experimental binding validation is a required next step before biological conclusions can be drawn. Second, the rigid-body docking approach (PatchDock/FireDock) does not capture receptor flexibility or carbohydrate-mediated adsorption; more recent multimeric docking tools such as AlphaFold3 or HADDOCK may improve accuracy in future iterations. Third, the *Listeria* dataset remains underrepresented relative to the *Salmonella* and *Escherichia* cohorts, representing a critical bottleneck that future database expansions must address. Finally, while the pipeline strategically integrates established tools rather than pioneering novel computational algorithms, its primary merit lies in optimising workflow interoperability and open-science accessibility rather than advancing raw methodological novelty.

Collectively, these findings challenge the assumption that extensive RBP sequence diversity necessarily precludes reliable computational annotation at scale: by combining protein language models with structural validation, high-confidence candidates can be prioritised even for underrepresented taxa such as *Listeria* phages. Future work should extend this framework to additional priority foodborne and clinical pathogens, and incorporate flexible or multimeric docking approaches (e.g., AlphaFold3, HADDOCK) to better capture carbohydrate-mediated adsorption interfaces missed by rigid-body docking. Experimental validation should prioritise two candidates in particular: binding assays for YDV08491.1, given its favourable predicted interaction with OmpF, and recombinant expression of YDV14336.1, given its direct structural and sequence homology to a crystallographically resolved *Salmonella* adsorption protein. Confirmation of host-binding activity for these candidates would represent a critical step toward their integration into phage-based biosensor platforms for foodborne pathogen detection.

## Materials and Methods

### Phage Genome Dataset Assembly

To build a representative dataset of phages relevant to food safety, complete genome sequences (as classified in NCBI Nucleotide) targeting *Salmonella* spp., *Escherichia coli*, and *Listeria monocytogenes* were retrieved from the NCBI Nucleotide database in April 2026. The initial dataset comprised 260 genomes: 100 *Salmonella* phages, 100 *Escherichia* phages, and 60 *Listeria* phages, the latter reflecting its status as a regulated pathogen under EU Regulation 2073/2005 and its relevance in dairy and ready-to-eat food matrices. These three genera were selected as representative priority pathogens across the food safety regulatory framework, spanning both Gram-negative (*Salmonella*, *Escherichia)* and Gram-positive (*Listeria*) foodborne threats. All code, datasets and predicted structures are publicly available, supporting reproducible development of phage-derived recognition elements for food safety applications github.com/XiomaraDRC/phage-rbp-foodborne-pathogens.

### Quality Filtering

A minimum genome length of 15,000 bp was applied to exclude incomplete or fragmented sequences, consistent with thresholds used for small-genome phages including *Podoviridae* and *Microviridae* families [13]. A minimum of 20 annotated coding sequences (CDS) was additionally required to ensure sufficient genomic content for downstream RBP prediction. These thresholds were defined based on established parameters in phage genomics literature [14].

### Protein Extraction

All annotated coding sequences (CDS) were extracted from the filtered genomes using Biopython. Furthermore, to ensure only translatable sequences were included in downstream analyses, CDS features lacking a translation qualifier were discarded.

### Receptor-Binding Protein Prediction

RBP candidates were identified using PhageRBPdetect v3 [10], a deep learning framework built on a fine-tuned ESM-2 protein language model. Predictions were executed *via* the Google Co-lab implementation with GPU acceleration, which enabled processing of the full protein dataset without local computational infrastructure as a practical consideration for research groups in resource-limited settings. Prediction thresholds of 0.80, 0.90, and 0.95 were evaluated. A cutoff of >0.90 was selected to define high-confidence RBP candidates, balancing false-positive reduction (Table 1).

### Structural Prediction of High-Confidence RBP Candidates

Three-dimensional structures of high-confidence RBP candidates were predicted using the ESMFold API. Prior to structural prediction, redundancy in the dataset was reduced by clustering sequences at 90% identity using CD-HIT (v4.8.1), yielding 306 unique representative sequences (102 *Salmonella*, 118 *Escherichia*, 56 *Listeria*). Sequences exceeding 400 amino acids were truncated to their C-terminal 400 residues, consistent with the known localisation of host-binding domains at the C-terminus of phage RBPs [5]. Of the 306 input sequences, 276 structures were successfully predicted (102 *Salmonella*, 118 *Escherichia*, 56 *Listeria*), corresponding to a 90.2% success rate. For Foldseek structural validation, the top five predicted structures per pathogen group were selected by ranked PhageRBPdetect confidence score among structures meeting the pLDDT ≥0.85 quality threshold, yielding 15 candidates submitted to structural homology search against PDB100.

### Protein–Protein Docking of Top RBP Candidates Against Host Receptors

To explore the binding potential of predicted RBP candidates against host surface receptors, exploratory protein–protein docking was performed using experimentally resolved receptor structures retrieved from the RCSB Protein Data Bank: OmpC from *Salmonella* spp. (PDB: 2J1N), OmpF from *Escherichia coli* (PDB: 2ZFG), and a wall teichoic acid-binding protein from *Listeria monocytogenes* (PDB: 4UDR), selected based on their established roles in phage adsorption [15]. The top-ranked RBP candidate per pathogen group, by PhageRBPdetect confidence score, was submitted to PatchDock (v1.3; clustering RMSD: 4.0 Å) for rigid-body geometric docking against its corresponding receptor. For the *E. coli*-targeting candidate (YDV08491.1), OmpF was selected specifically for its high abundance and structural characterisation as a primary porin exploited during initial phage adsorption. The highest-scoring solution per pair was refined using FireDock with medium refinement parameters and 10 output structures. Binding quality was assessed through three complementary metrics: PatchDock geometric score, interface area, and Atomic Contact Energy (ACE; kcal/mol). These docking analyses are treated as exploratory and hypothesis-generating; results are not presented as evidence of confirmed binding in the absence of experimental validation.

## Acknowledgments

This research received no specific grant from any funding agency in the public, commercial, or not-for-profit sectors. The authors are grateful to the Instituto de Investigación y Desarrollo de Procesos Químicos (IIDEPROQ) at Universidad Mayor de San Andrés (UMSA) for providing institutional support.

## Data and code availability

All scripts, processed datasets, accession lists, predicted structures, and reproducible workflows are publicly available at https://github.com/XiomaraDRC/phage-rbp-foodborne-pathogens. Raw phage genome sequences were retrieved from NCBI Nucleotide (accession numbers listed in Supplementary Table S1). No newly generated experimental data is associated with this study.

---

**Figure 1.**
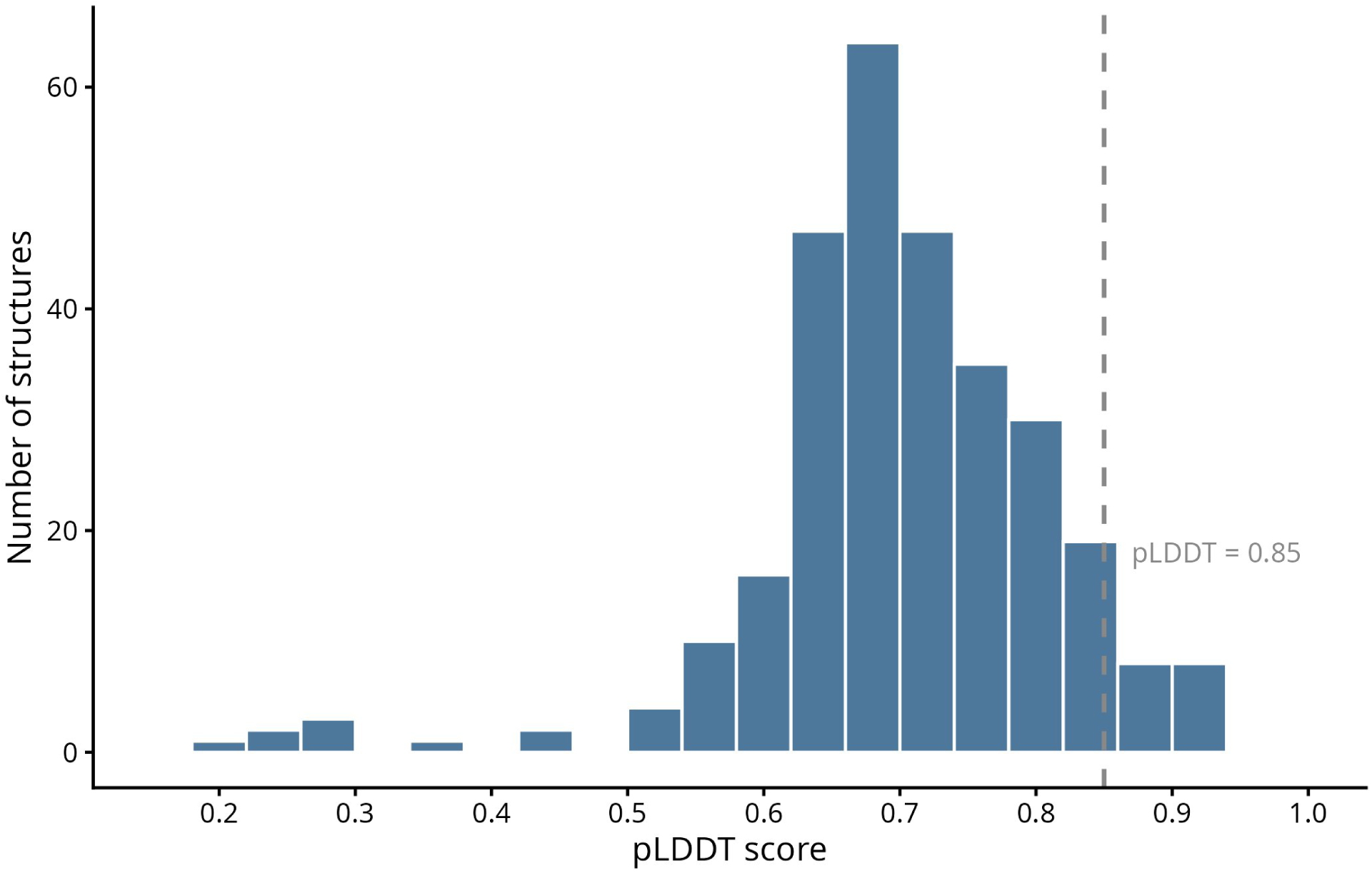
pLDDT score distribution of ESMFold-predicted RBP structures (n = 276 unique structures). Dashed line indicates the pLDDT ≥0.85 selection threshold applied for downstream Foldseek structural validation.

**Figure 2.**
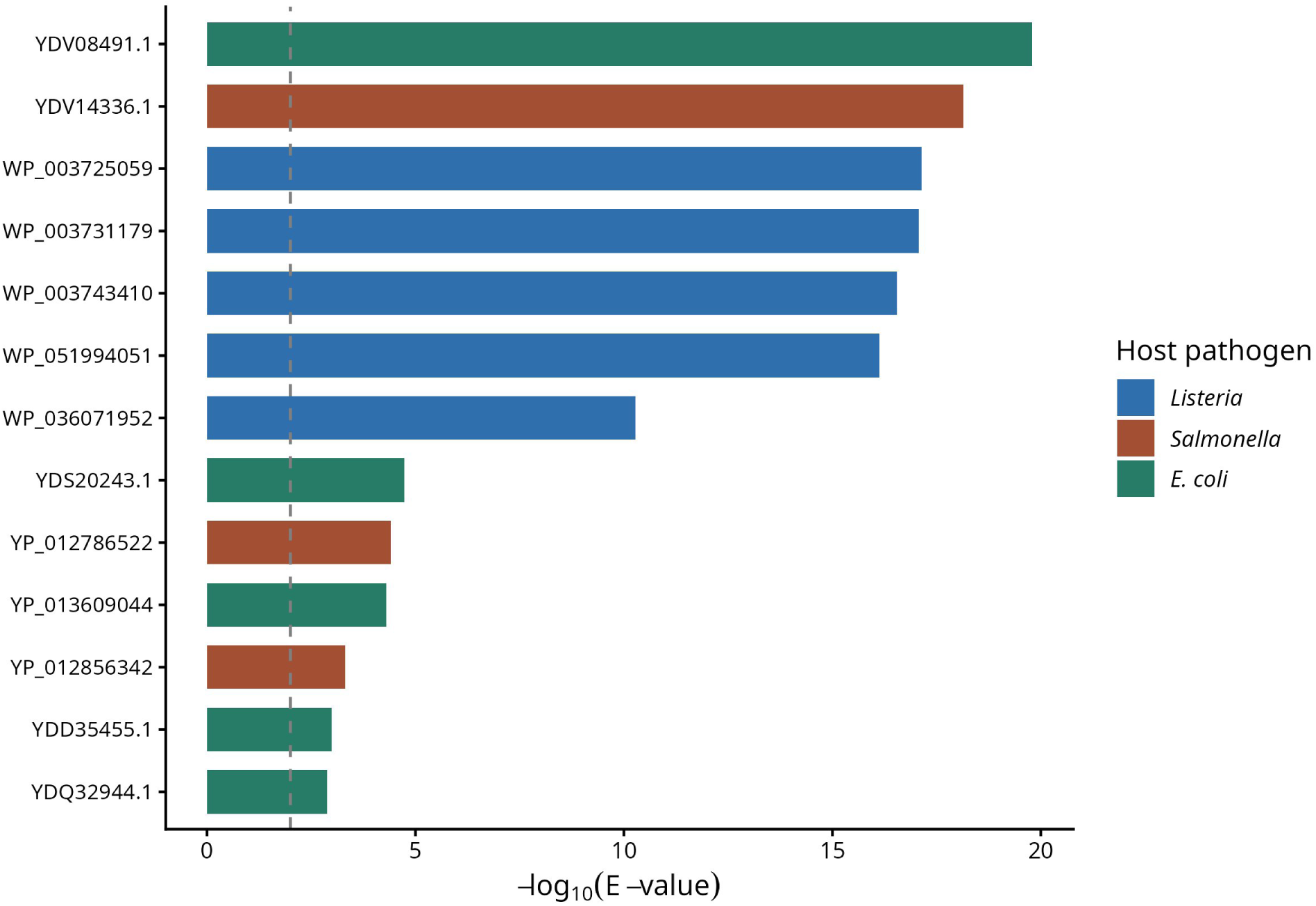
Foldseek structural validation of the top 13 RBP candidates against PDB100. Bars represent −log_10_(E-value) for the top hit per candidate. All candidates returned probability = 1.0. Colours indicate host pathogen group.

**Figure 3.**
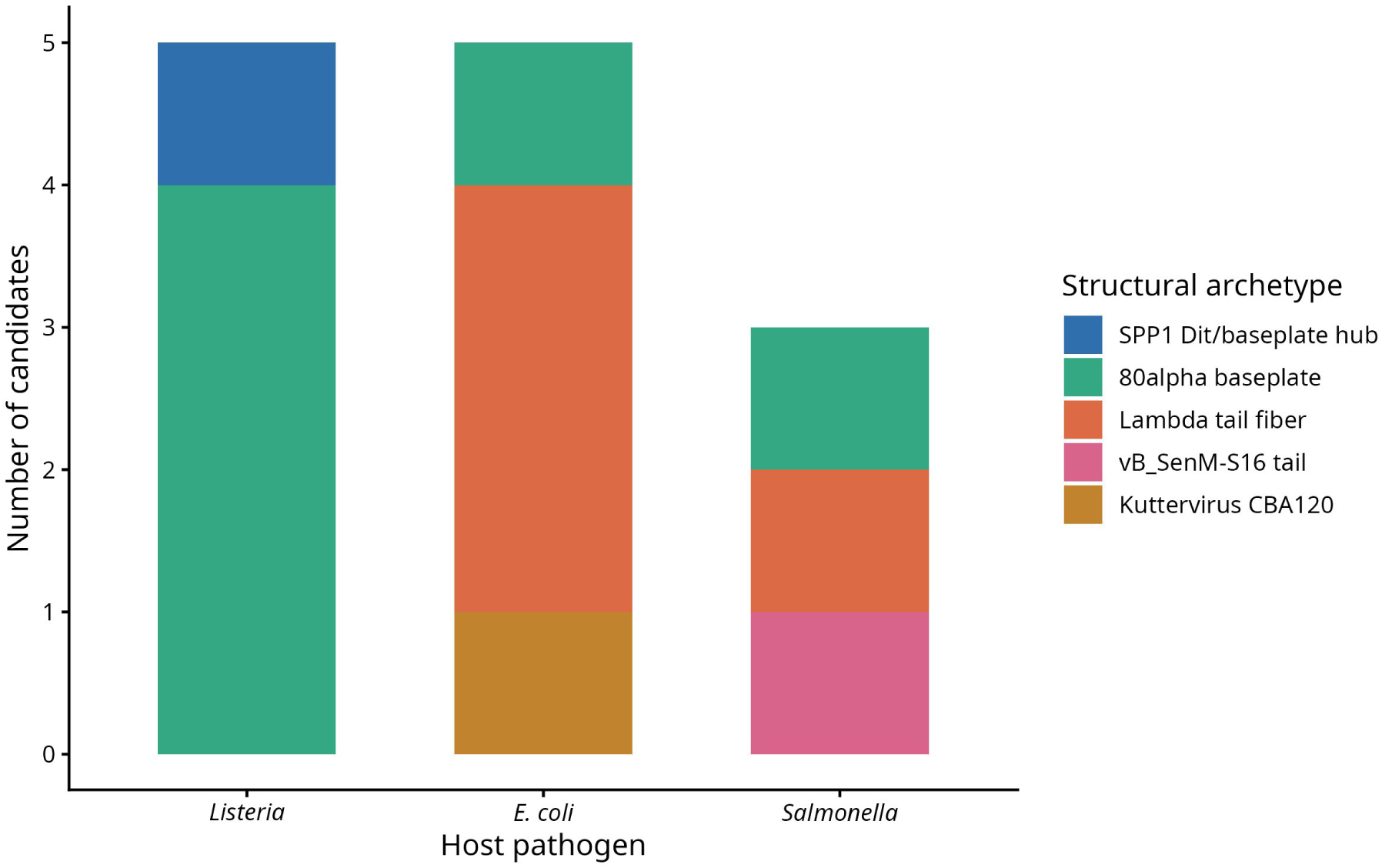
Distribution of structural archetypes among the 13 validated RBP candidates by host pathogen group, based on top Foldseek hits against PDB100.

**Figure 4.**
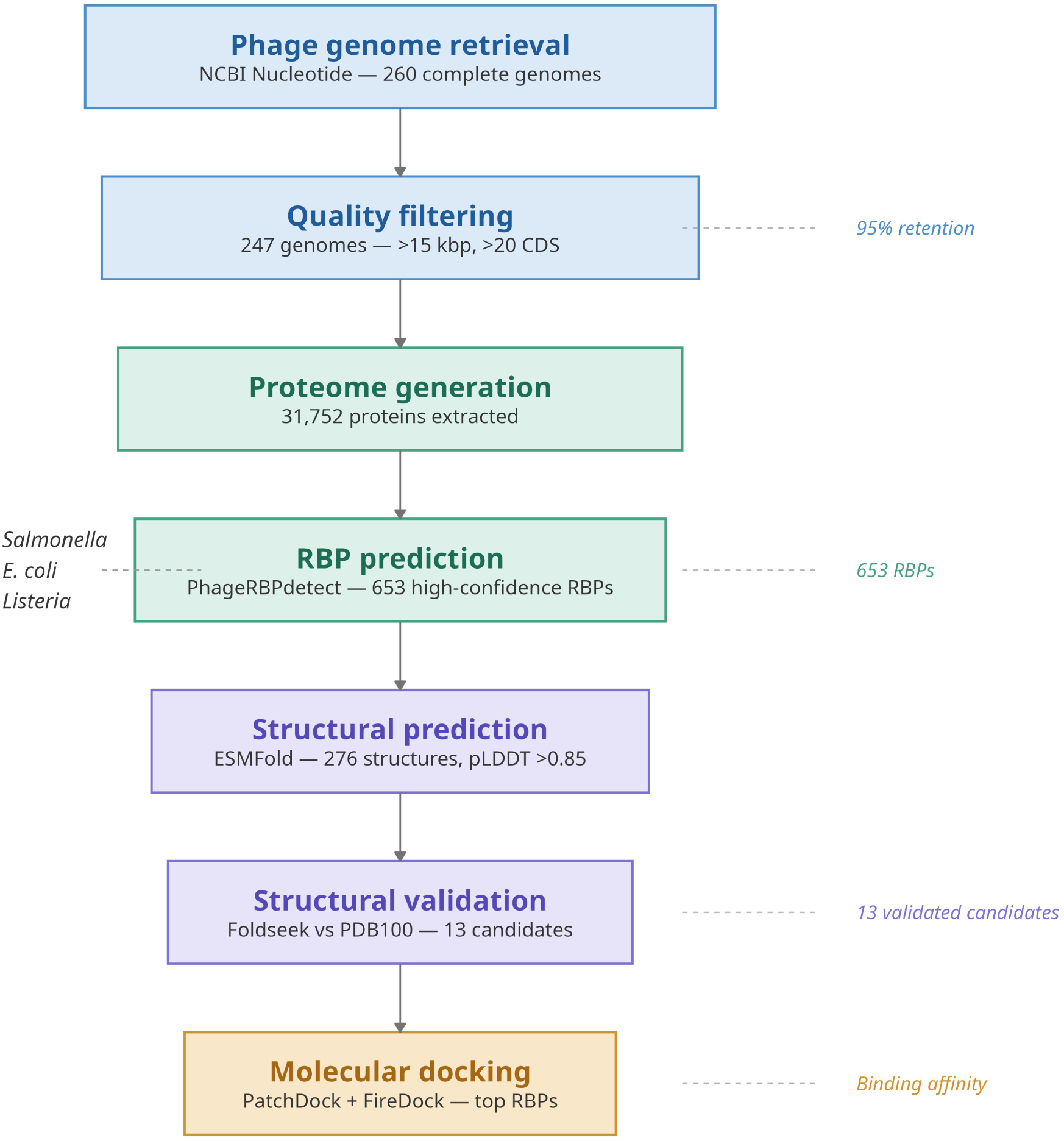
Overview of the computational pipeline for phage RBP identification, from genome retrieval through molecular docking validation.

**Figure 5.**
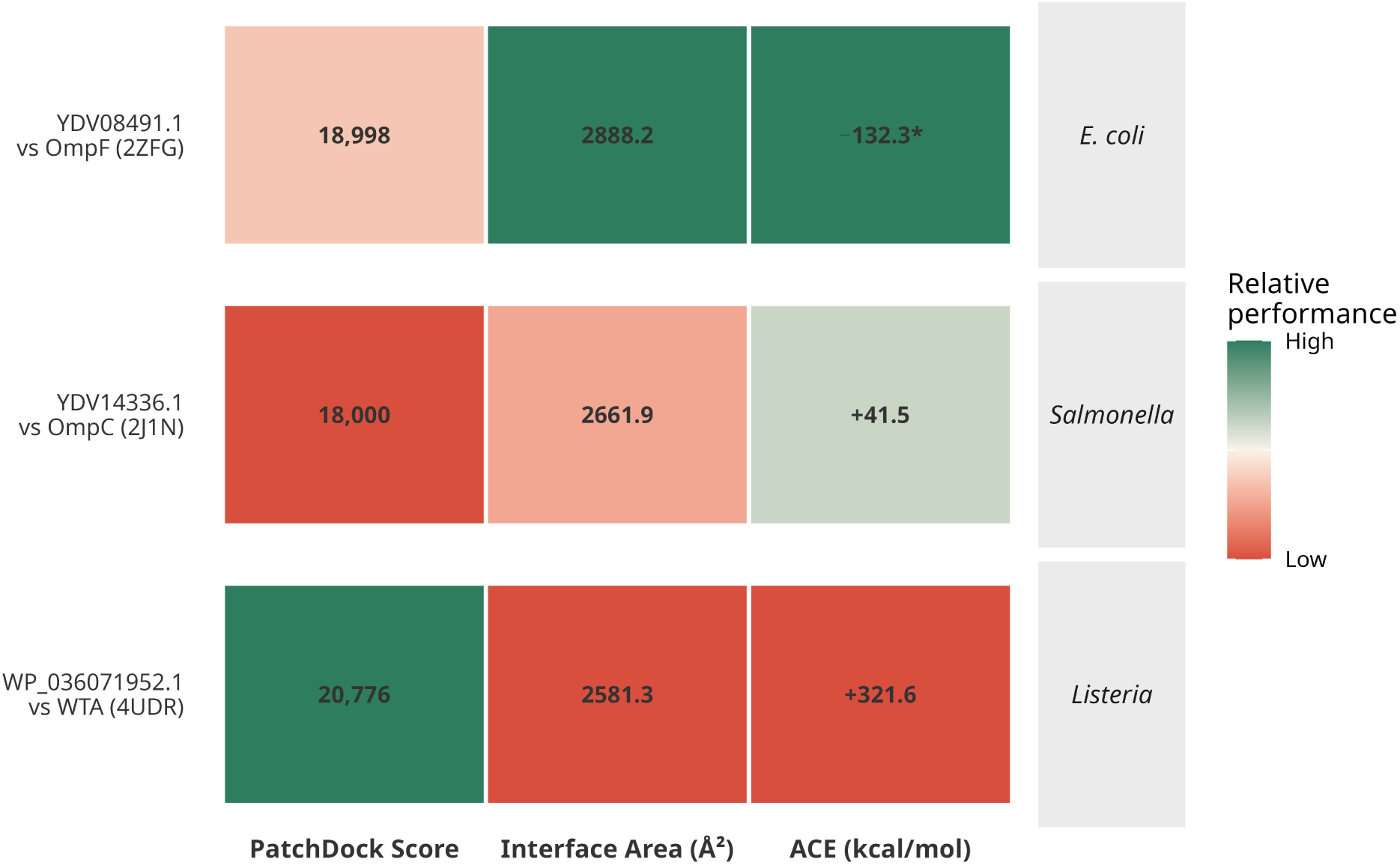
Heatmap of protein–protein docking results for the top RBP candidate per pathogen group against cognate host surface receptors. Colour intensity reflects relative performance per metric (green = high, red = low). For ACE, lower (more negative) values indicate energetically favourable binding; asterisk (*) marks the only candidate with a negative ACE value (YDV08491.1 vs OmpF; ACE = −132.3 kcal/mol).

